# An animal society based on kin competition, not kin cooperation

**DOI:** 10.1101/854802

**Authors:** Jessica L. Vickruck, Miriam H. Richards

## Abstract

Animals respond to competition among kin for critical breeding resources in two ways: avoidance of direct fitness costs via dispersal of siblings to breed separately, and formation of kin-based societies in which subordinates offset direct fitness costs of breeding competition via altruism and increased indirect fitness. Here we provide the first evidence that kin competition can promote the evolution of societies based on non-kin cooperation. For eastern carpenter bees, nests are a critical breeding resource in perpetually short supply, leading to strong competition among females. Observations of individually marked and genotyped females demonstrate that sisters disperse from their natal nests to join social groups of nonrelatives. By forming social groups of non-kin, females increase their chances of successful reproduction, while avoiding the indirect fitness cost of competition among sisters.

**One Sentence Summary:** We describe the first known example of an animal society based on avoidance of kin competition rather than on promotion of kin cooperation.

## Main Text

Ultimately, competition for crucial resources linked to reproduction, such as food or breeding sites, shapes the evolution of social behaviour in animals. One way for individuals to better extract or protect resources is to do so jointly, in cooperation with other individuals; cooperative and helping behaviours are a major reason why groups of individuals can have higher per-capita fitness than solitary individuals (*1*). However, individuals living in groups do not escape competition: often they are vulnerable to within-group competition for breeding (*2–4*) resources, competition that may be especially pronounced for limited resources, such as nests. While it is possible for group members to share resources equitably, most animal societies are hierarchical, rather than egalitarian (*1*). Moreover, most animal societies are kin-based (*1*). In hierarchical, kin-based societies, subordinates can respond to exploitation and unequal sharing of resources by resisting or avoiding selfish exploiters, for instance, by dispersing to new locations, and avoiding breeding competition with close kin (*5*). Alternatively, subordinates cheated by relatives can potentially make the best of a bad situation or even prosper through the indirect benefits of helping their reproducing relatives to rear more offspring (*2, 3, 6, 7*). Considerable research has built a robust theoretical framework for understanding the consequences of kin competition (*3, 4*). Competitive interactions tend to lower the direct fitness of competitors, especially for those with lower competitive ability. When competitors are related, competition additionally lowers indirect fitness, so the inclusive fitness consequences of are predicted to be especially severe. One way of avoiding the negative consequences of kin competition, is for offspring to disperse prior to breeding, so as to avoid harming relatives that carry the same genes, and most empirical studies have addressed how kin competition influences dispersal (e.g. (*5*). However, dispersal also lowers opportunities for cooperative interactions that might increase the ability of cooperators to secure resources for their group, and results in decreasing population viscosity and relatedness among cooperators, which decreases the inclusive fitness payoffs of cooperation and altruism (*7*). Some theory predicts that the consequences of competition for resources within kin groups can be severe enough to “totally negate” the inclusive fitness benefits of kin cooperation (*7, 8*). Nevertheless, the existence of numerous animal societies based on kin cooperation in both vertebrates and invertebrates (*1*) suggests they evolved to overcome the disadvantages of kin competition (*9*). Here we describe the social behaviour of a facultatively social bee in which group formation and social structure balances cooperation with competition for breeding resources, providing the first known example of an animal society based on avoidance of kin competition rather than promotion of kin cooperation.

Facultatively social animals are ideal models for examining the ecological conditions that promote group or solitary living, because they exemplify the costs and benefits of solitary versus social living under varying ecological conditions (*10, 11*). In eastern carpenter bees, *Xylocopa virginica*, decisions to nest solitarily or socially (in groups) are linked to competition for access to nests, which are critical breeding resources that are costly to construct and occupied by successive generations of females, sometimes for decades (*12*). Most females breed in old nests, remaining in their natal nests or relocating to neighbouring nests containing groups of unrelated females; much less frequently, a female initiates construction of a new nest. When nesting in groups, female carpenter bees form linear reproductive queues in which the dominant female monopolizes reproduction, while subordinates wait for opportunities to replace her (*13, 14*). Group size is usually 2-4 females, but may be as high as 8; refs (*14, 15*) and below. Thus, females experience intense breeding competition at the population level for access to nest sites, and within social groups for egg-laying opportunities. Since only one female at a time can lay eggs, the average inclusive fitness of related females that remain together in their natal nest, will be lower than if only one remains in the natal nest, while the others disperse to breed in separate nests.

Here we demonstrate that in spring, intense competition among females for nest sites increases both the frequency of social group formation (nest-sharing) and the intensity of within-group competition for breeding opportunities. We then demonstrate that females avoid competition with relatives by dispersing from their natal groups to preferentially join breeding groups composed of non-kin. While a shortage of nest sites imposes unavoidable competition on females for this critical resource, essentially females choose to compete with non-relatives, avoiding the indirect fitness costs of competing with relatives.

Inter-individual resource competition should be most severe when critical resources are relatively scarce or when population density is high; competition for limited resources such as breeding sites should be particularly severe (*9*). We predicted that high population density would result in more frequent social nesting and larger average colony size. Since larger colony size would mean lengthened reproductive queues, we also predicted that in high density, bees would more frequently attempt to improve their social position by leaving their natal nests to join new colonies, either nearby or by leaving the population altogether to breed elsewhere. To test these predictions, we carried out detailed behavioural observations of female carpenter bees at several nesting aggregations in wooden bridges in a park near Brock University in southern Ontario, Canada (Supplementary Figure 1). This location is close to the northern edge of the species’ range, with shorter flight seasons and much more severe winters than in southern locations where it has been previously studied (*15*). Eastern carpenter bees overwinter as adults in their natal nests, with females awaking from hibernation in early spring (April in southern Ontario). In early spring, we trapped and individually marked all bees with enamel paint on their first venture outside their natal nests. The activity patterns of eastern carpenter bees are unusual for social bees, with two foraging flight periods (*13*). The first phase is a nestmate provisioning phase (NMP) in which dominant females bring pollen back to the nest to feed to adult nestmates; this reinforces linear reproductive queues established prior to the onset of NMP. The second phase is a brood provisioning phase (BPP) in which the dominant female provisions brood cells in which only she lays eggs. If the dominant female dies, disappears or becomes moribund, the next female in line replaces her as the principal forager and egg-layer (*13, 14*). During NMP and as late as BPP, secondary females often leave their natal nests to join neighbouring nests in the same aggregation, other aggregations, or leave the area altogether to nest elsewhere. Brood provisioning ceases in early July, after which adult females remain inside the nests with developing brood. When brood eclose as adults at the end of the summer, they may be fed by older adult females that leave the nests during a brief, late summer foraging phase (late August or early September in southern Ontario). The newly eclosed adults, both male and female, remain in their natal nests throughout the winter until the following spring. Older females occasionally survive through a second winter, but virtually always die or disappear early in the NMP.

In general, competition for resources is higher when population densities increase or when resources become scarcer (*5, 9*). For large carpenter bees in general, and eastern carpenter bees in particular, nests are a critical breeding resource, costly to produce and in perpetually short supply. Eastern carpenter bees forage on a wide variety of blossom types, but their nests are almost always found in structures built of milled lumber, especially pine and spruce (*16*); thus they are foraging generalists but nesting substrate specialists (*17*). They are strongly philopatric and nesting aggregations persist for years or even decades, as successive generations of females reuse nests (*12, 13, 15*). Nests are very costly to construct; a female that initiates nest construction may take up to a week to construct a nest with a single tunnel, during a BPP that lasts only 3-6 weeks (note that nests are never founded jointly). As a result, most females attempt to breed in their natal nest or relocate to existing nests close by, usually in the same aggregation (*18*).

Eastern carpenter bees are facultatively social, most females nesting in groups, while a few females nest solitarily (*19*). Under crowded, high density conditions, female bees should be more likely to share nests. We examined bee nesting behaviour during 2012 when population density was high, and 2013 when population density was low. As predicted, group size was strongly associated with population density (Figure 1). In high density conditions (2012), 96% of overwintered females nested socially (in groups) and average group size was significantly higher than in 2013 when population density was much lower and only 70% of females nested socially (Table 1). In 2012, females were more likely to relocate from their natal nest to another nest, and proportionately more females left the aggregation to nest elsewhere. Moreover, in 2012, 9 females initiated new nests, a rate of new nest construction unprecedented at this location over 7 years of observations (2011-2013, 2016-2019); no new nests were initiated in 2013. In every instance of nest initiation observed at our study sites (several dozen examples), a single female excavated the nest entrance and the first sections of tunnel by herself. Thus, nest initiation is a solitary activity, although additional females frequently join the new nest within a day or two of nest entrance completion. The significant differences in group size between 2012 and 2013 demonstrate density dependence of social group formation, with strong competition for nest sites inducing higher rates of dispersal, higher rates of nest construction, higher rates of social group formation, and increased social group size.

**Figure 1.**
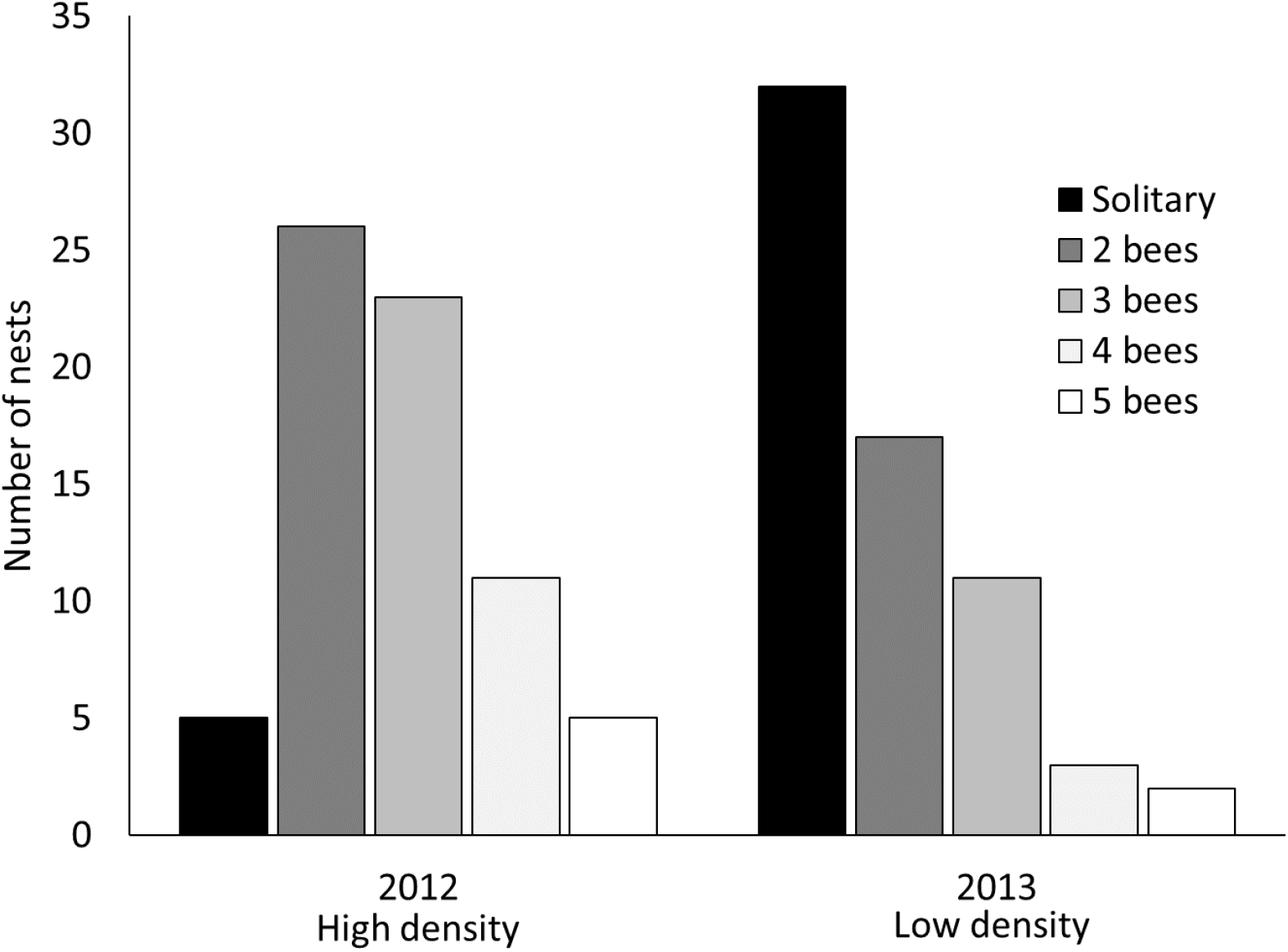
Variation in group size during the brood provisioning phase of the colony cycle in response to high (2012) and low (2013) population density. In low density, significantly more females nested solitarily (Table 1, P<0.00001, also see Supplementary Table 1). In both 2012 and 2013, the maximum colony size during the brood provisioning phase was 5 females.

**Table 1.** Female competition for nest sites varies with population density. Adult females were caught and marked when they first emerged from hibernation in the spring, so the number of females marked each year is an accurate measure of population density. Nesting location, number of occupied nests, and colony size were estimated during the brood provisioning phase in June and July, which lasted 31 days in 2012 and 32 days in 2013.

|  | Year |  |  |
| --- | --- | --- | --- |
|  | 2012<br>High density | 2013<br>Low density | 2012 vs. 2013 |
| Females marked | 189 | 101 |  |
| Nesting location <sup>1</sup> |  |  |  |
| Natal nest (residents) | 74 (46%) | 65 (64%) |  |
| Different nest (transients) | 34 (21%) | 17 (17%) |  |
| Disappeared | 52 (33%) | 19 (19%) | X<sup>2</sup>= 8.70, d.f.= 2, P=0.01 |
| Number of nests <sup>2</sup> | 70 | 65 |  |
| Solitary | 5 (7%) | 30 (46%) |  |
| Social | 65 (93%) | 35 (54%) | X<sup>2</sup>= 26.35, d.f.= 1, P<0.00001 |
| Colony size during the brood provisioning period (females per nest) | 2.79 ± 1.03 | 1.86 ± 1.06 | Mann Whitney U= 1167, P< 0.0001 |

<sup>1</sup> In 2012, 29 females in were caught flying through the nesting aggregation, so natal nest was unknown, and they could not be assigned resident or transient status.
<sup>2</sup> Newly constructed nests were included in the social nest category; 9 new nests, each of which contained 2 females, were constructed in 2012 when competition for nest sites was more severe, and no new nests were constructed in 2013.

While population-level competition for available nest sites is the critical factor driving group formation in *X. virginica*, it also intensifies within-group competition for breeding opportunities. In eastern carpenter bees, only the dominant female in a social group forages and lays eggs, whereas subordinates queue for the opportunity to replace her when she dies or becomes moribund (*14*). Whereas group size was significantly higher in 2012 than in 2013, the duration of the brood provisioning phase of the colony cycle was the same (Table 1). As a result, fewer subordinates would have achieved egg-layer status under high density, compared to low density. In a previous study (*13*), primary females (rank 1) provisioned the most brood, but often died before the end of the BPP. As a result, secondary females in ranks 2 and 3 often achieved breeder status partway through the brood-provisioning season, although females in ranks 4 to 6 never did. Moreover, in our study population, primary and second females virtually never survive to a second breeding season (*14*), so their fitness is completely predicated upon raising brood in the current breeding season. Thus, the significantly longer reproductive queues of 2012 would have led to significantly lower average direct fitness of subordinate females that year.

In virtually all social insects, groups are composed largely of kin; although recent studies have highlighted the fact that colonies often contain unrelated individuals, these always comprise a minority of the colony (<15% of colony members). Cooperative breeding can enhance both direct and indirect fitness when cooperators are related, whereas unrelated individuals can only gain direct fitness. To determine whether eastern carpenter bee breeding groups are composed of kin or non-kin, we genotyped female nestmates at nine microsatellite loci (*20*), then used Kingroup V2 (*21*) to identify nestmates likely to be full sisters (*22*). We examined groups at three successive time points: in late winter when females were still in their natal nests; in spring during the nestmate provisioning period, when females establish reproductive queues in social nests; and in summer during the brood provisioning period when dominant females provision their brood and lay eggs. The proportion of full sisters in colonies declined significantly from winter through spring to summer (Figure 2, Supplementary Table 2), suggesting that sibships were broken up as females dispersed to nests in spring. Relatively high relatedness among cohabitating females in winter, suggests that many females could have formed kin groups in spring. We therefore identified all possible full sisters for each genotyped female in the population and investigated whether they nested together or apart (Figure 3). Of 266 genotyped females that were still alive at the time of brood provisioning and egg-laying, only 30 sisters (11%) nested together, 178 sisters (67%) nested apart (in different nests), and 58 (22%) did not have a full sister in the population (there was no difference between years). The proportion of sisters nesting together in summer was not significantly different from expectation under a null hypothesis in which females were randomly distributed among nests (Supplementary Figure 2). Overall, these results indicate that siblings overwintered together but kin relationships were broken up in spring as many females relocated to new nests. This suggests that dispersal from the natal nest is a mechanism by which females solve the problem of severe kin competition for breeding opportunities in their natal nests. Once they leave their natal kin groups, females still face strong competition for nest sites, which forces them to share nests and join social breeding groups. By doing this, they incur the direct fitness costs of breeding competition with non-relatives, but avoid additional, indirect fitness costs of breeding competition with relatives.

**Figure 2.**
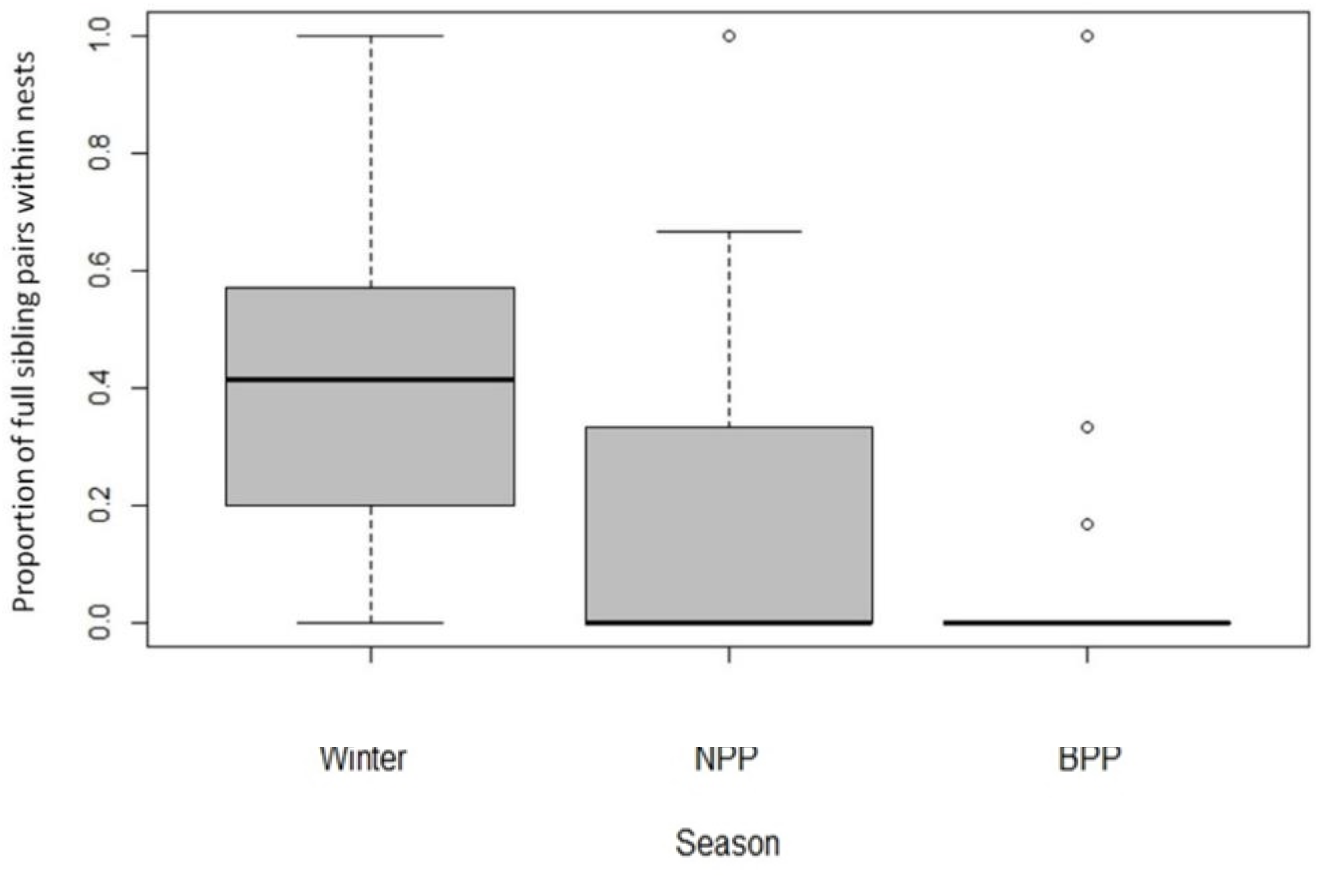
Decline in kinship among social nestmates prior to the formation of breeding associations. Winter associations represent natal nestmates, since bees overwinter in their natal nests. The proportion of nest mate pairs that were full sisters decreased significantly from winter, through the spring nestmate provisioning phase (NPP) to the summer brood provisioning phase (BPP) (Kruskal-Wallis X^2^=13.01, d.f.=2, P=0.001). Boxes-and-whiskers represent median and quartile ranges, while open circles represent outliers.

**Figure 3.**
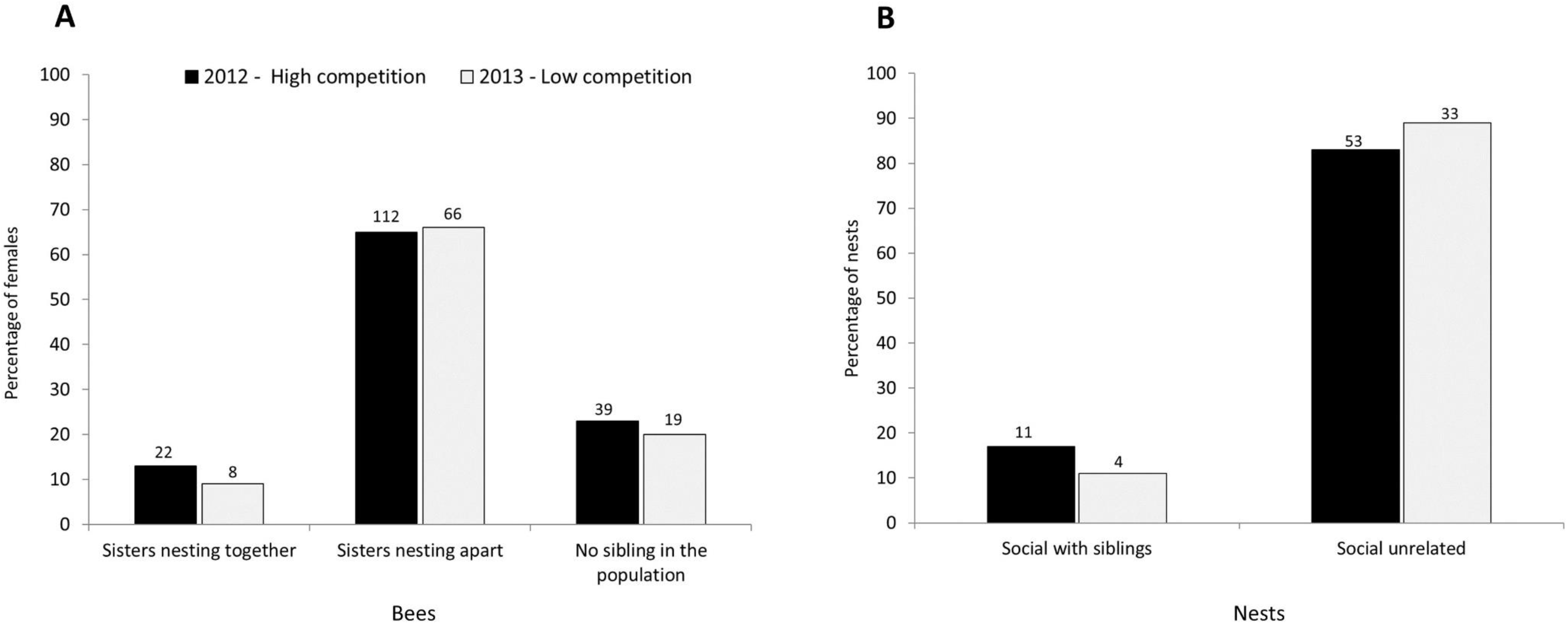
Evidence for avoidance of kin competition by adult females during the brood provisioning period. **(A)** Although most females had at least one full sister in the population, most females nested apart from their sisters, rather than remaining together in the natal nest or dispersing together to a new nest. The proportions of sisters nesting together or apart did not differ between 2012 and 2013 (X^2^=1.38, d.f.=2, P=0.50), suggesting that changes in population density did not influence the relatedness structure of colonies. **(B)** The proportion of social colonies that contained full sisters was also similar in 2012 and 2013 (Fisher’s exact test, P=0.48). Numbers above the bars represent the number of individuals **(A)** or nests **(B)**.

In most social insects, which live in kin groups, cooperative and helpful behaviours are directed at relatives. Eastern carpenter bees nesting in non-kin groups also provide evidence for cooperation and helping behaviours. Females are more tolerant toward nestmates and more aggressive to non-nestmates (*18, 23*), and both dominant and subordinate females have been observed to guard nest entrances against conspecific intruders. The most dramatic example of cooperation occurs in early spring, prior to the onset of egg-laying and brood provisioning, when dominant females forage for pollen that they feed to subordinate nestmates, especially “tertiaries” that rarely leave the nest (*19, 23*). Such observations demonstrate that cooperative behaviour can evolve in societies of non-kin, apparently via selection on the direct fitness of cooperators, rather than through the indirect fitness of helpers, as in the vast majority of social animals (*7*).

Although this is the first example of an animal society based on avoidance of kin competition, there are intriguing hints in other social insects that nonkin-based sociality may often have been overlooked. To start with, many carpenter bees are facultatively social, nesting either solitarily or in small groups, sometimes as few as two females, in which only one female lays eggs, while other females await their turn to become the primary reproductive (*24*). Accumulating behavioural evidence suggests that some euglossine bees are facultatively social, forming social groups that resemble reproductive queues and which may include nonkin. Recent phylogenetic evidence suggests that societies based on reproductive queues in which subordinates wait for their own reproductive opportunities could represent an early stage in evolutionary transitions to caste-based sociality, in which subordinates, sacrifice their own reproduction to aid kin (*25, 26*). Indeed, it is now well known that many eusocial insect species, in which societies are based primarily on kin-based, reproductive altruism, a substantial minority of females exhibit reproductive strategies that may represent avoidance of kin competition. In recent years, considerable evidence has accumulated that subordinate individuals, variously known as “drifters”, “aliens”, or “joiners” leave their natal colonies and join unrelated colonies (*27, 28*). Some wait for opportunities to inherit the role of dominant egg-layer (the ‘queen’ role in eusocial colonies). Whether such joiners actively avoid moving to nests with relatives should be investigated. In many ants, unrelated gynes cooperate to initiate new nests, but once the first workers emerge, they aggressively and often violently, compete for dominance, with a single female becoming the colony’s egg-laying queen (*29*).

The ubiquity of within-group competition likely explains why most animal societies, and virtually all social insect societies, are based on reproductive altruism, in which subordinate helpers sacrifice some or all of their own reproduction to enhance that of related reproductives (*30*). Reproductive altruism by subordinates is one solution to the problem of intense kin competition within groups, and judging by the repeated evolution of eusociality and cooperative breeding, is a highly successful solution (*31*). Eastern carpenter bees demonstrate alternative solutions to the problem; when the availability of nests is high, females can avoid kin competition by nesting solitarily, and when availability is low, they avoid kin competition by nesting in non-kin groups.

## Acknowledgments

We sincerely thank Nigel Raine, Liette Vasseur, Glenn Tattersall, and especially Graham Thompson, for their comments and input on this research. This research was funded by a Natural Sciences and Engineering Research Council (NSERC) of Canada Discovery Grant to MR and an NSERC Post-Graduate Scholarship to JV, as well as internal funding and resources from Brock University.

## Data and materials availability

All data are available in the main paper or in the Supplementary Materials

## List of Supplementary Materials

**Materials and Methods**

**Supplementary Table 1.** Shift in colony size distribution during the brood provisioning phase of the colony cycle, from high population density in 2012 to low population density in 2013.

**Supplementary Table 2.** Decline in mean relatedness among nestmates from the late winter hibernation phase to nestmate provisioning phase (NPP) to brood provisioning phase (BPP).

**Supplementary Table 3.** Microsatellite genotypes for all Xylocopa virginica females captured in 2012 and 2013 that were used for inferring sibling relationships and relatedness.

**Supplementary Figure 1.** Map of sample sites.

**Supplementary Figure 2.** Simulations to assess the number of sibships that would be observed if genotyped females were randomly distributed among available social colonies in and 2013.

