## Supplemental tables, figures, data for "An animal society based on kin competition, not kin cooperation"

### SUPPLEMENTARY MATERIALS

#### Methods

*Description of field sites*

We studied five nesting aggregations of *Xylocopa virginica*, each one located in a wooden bridge at the Glenridge Quarry Naturalization Site (GQNS), in St. Catharines, Ontario, Canada (43.122, -79.236 decimal degrees). Each bridge was home to 10-22 nests. Bridges were constructed in 2003 and were available for the bees to use as nesting substrate beginning the spring of 2004. Eastern carpenter bees often reuse nests for many years. The term 'new nest' refers to nests that were constructed in the current observation year, while 'old nest' refers to nests that are being reused from previous years.

X. virginica *colony cycle*

In southern Ontario, bees emerge from hibernation around late April or early May. *Xylocopa virginica* females have two distinct foraging phases (*1*). The nestmate provisioning phase (NPP) takes place just after spring emergence and females bring back pollen to the nest to feed other adult conspecifics rather than to provision brood (*1*). The NPP is also the time when females can disperse from the nest in which they overwintered. Females were categorized as resident or transient based on behavioural observations over the course of the foraging season. Resident females did not disperse and were only ever seen in one nest, while transient females dispersed from the nest in which they overwintered and were observed in more than one nest across the season. The NPP is followed by the brood provisioning phase (BPP) where pollen brought back to the nest is used to create the large pollen balls on which females lay their eggs. The BPP generally ends during the second week of July; thereafter, adult females rarely venture outside their nests. Once brood have been provisioned females remain in the nest to guard developing offspring from predators and parasites. Offspring eclose from late-July to August, with both female and male offspring remaining in their natal nests to hibernate until the following spring. In southern Ontario, most females live one year, breeding in their first spring and summer following eclosion, rarely surviving a second hibernation. In more than 7 years of intensive behavioural observations, no female has ever been observed to forage in two years, so lifetime reproductive success is equivalent to the number of brood produced in one breeding season.

*Bee handling and observations*

Bees were captured at nest entrances using cup traps (*2*). Cup traps are medium sized plastic cups that have a small hole cut out of the bottom while the top is covered with parafilm. The end with the small hole is then secured over the nest entrance with Velcro. Bees leave the nest but become trapped in the cup. Unmarked females were immediately placed on ice for approximately 10 minutes to allow for individual marking and measurements to be taken. Each female was individually marked on the thorax with a unique two-colour combination using enamel model paint. Bees were measured across the widest part of their heads (head width) to use as a proxy for body size comparisons. At this time the last tarsus of the left mesothoracic leg was removed and placed in chilled 100% redistilled ethanol for genotyping at a later date. Marked and measured bees were placed back outside of their nests to warm up and resume activity. There was no indication that marking bees disrupted their ability to locate their nest (*1*).

Foraging observations took place for 8h periods (8:00 to 16:00) during the nestmate provisioning phase on days where there was no rain and the temperature was greater than 20 °C. Time and nest of departure as well as the bee's individual paint ID were recorded for each bee leaving the nest. Bees were then released and the trap was replaced over the nest entrance. When a bee returned to the nest, the trap was removed to allow her entry. The time of her return, the nest which she returned to, as well as whether or not she was carrying pollen was recorded. At the end of the day all cup traps were removed for the night. Observations ceased when an entire observation day passed (8h) without seeing a single pollen trip by a female bee, indicating that the brood provisioning phase was complete.

To quantify the relationships among individuals inside overwintering nests, nineteen nests were destructively sampled in March of 2012.  These nests were carefully planed open to expose overwintering bees. All individuals inside nests were measured, marked and had a tarsal sample taken using the same techniques as summer bees described above.

*Assigning nest status*

Nest status (social or solitary) was determined by observations across the entire season. Each time a female departed from the nest a small, flexible plastic transfer pipette was inserted in the nest entrance. If there was still a female present in the nest she would buzz, bite the pipette tip or block the entrance with her abdomen. The presence or absence of a guarding female was recorded and used to determine if the nest was social or solitary. Nests were classified as solitary if during the brood provisioning phase only one female was ever seen bringing pollen to the nest and a second bee was never observed guarding the nest entrance. Nests were classified as social if more than one female was recorded in the nest during the brood provisioning phase.

*Genetic analyses and relatedness calculations*

DNA extraction and genotyping procedures are described in (*3*). In 2012, 189 females from 71 nests were genotyped. In 2013, 101 females from 64 nests were genotyped. Sixteen females were excluded from analyses of relationships in 2012 and 8 in 2013 due to missing data at more than 2 loci. Microsatellite genotypes are provided in Supplementary Table 3.

Relatedness among female nestmates was calculated using the method described by (*4*) as implemented in the program Kingroup V2 (*5*). Kingroup V2 allowed us to differentiate which pairs of bees within nests were significantly more likely to be full sisters than unrelated pairs. Hymenoptera are haplodiploid (females are diploid while males are haploid) therefore full sisters inherit one of two maternal alleles and must inherit the single paternal allele. When comparing full sisters, this means that full sisters must share the paternal allele at all loci.

*Modelling relatedness distributions*

We used a randomization analysis to determine if the number of sisters observed nesting together in the population was different from the number of sisters that would be observed nesting together if females were randomly distributed into nests. To do this we assigned all females marked in either 2012 or 2013 to simulated nests at random. In each sample year, the number of nests as well as the size of the nest (the number of females recorded inside) was replicated exactly as was seen in the sample population. After females were randomly assorted into nests, we used Kingroup V2 (*5*) to determine how many full sister pairs were present in simulated nests, as well as how many simulated nests contained full siblings. We then repeated this procedure 100 times for both the 2012 and 2013 datasets.  Simulation results were used create distributions for the expected number of siblings in nests and the number of nests that contained full siblings given the bees in the population for both 2012 and 2013.  We then compared our observed values to the expected distributions of our randomization analysis to quantify the probability of our observations given the simulated data.

**Supplementary Table 1.** Shift in colony size distribution during the brood provisioning phase of the colony cycle, from high population density in 2012 to low population density in 2013. Census nests represent the subset of the population in which the number of females was most accurately estimated in a removal experiment.

| | **All nests** | | **Census nests** | |
|---|---|---|---|---|
| **Number of nests** | **2012** | **2013** | **2012** | **2013** |
| 1 (solitary) | 5 (7%) | 32 (49%) | 5 (11%) | 30 (63%) |
| 2 | 26 (37%) | 17 (26%) | 7 (16%) | 5 (10%) |
| 3 | 23 (33%) | 11 (17%) | 19 (43%) | 10 (20%) |
| 4 | 11 (16%) | 3 (5%) | 10 (23%) | 3 (6%) |
| **5** | 5 (7%) | 2 (3%) | 3 (7%) | 0 (0%) |
| **Total** | 70 (100%) | 65 (100%) | 44 (100%) | 48 (100%) |
| **Females per nest** | 2.79 $\pm$ 1.03 | 1.86 $\pm$ 1.06 | 2.98 $\pm$ 1.07 | 1.71 $\pm$ 1.21 |
| **Comparison between years (Mann Whitney U-test)** | U=1167, P<0.00001 | | U= 429.5, P<0.00001 | |

**Supplementary Table 2.** Decline in mean relatedness among nestmates from the late winter hibernation phase to nestmate provisioning phase (NPP) to brood provisioning phase (BPP). Values in brackets represent the number of nests. Queller-Goodnight estimates of average relatedness were calculated for all possible pairs of female nestmates within each colony, based on female genotypes at 9 microsatellite loci (Vickruck and Richards 2017). The proportion of full sisters is the proportion of all possible female pairs in each nest whose genotypes suggested that they were full sisters, using Kingroup V2 (Konovalov 2004). The median proportion of nest mate pairs that were full sisters decreased from winter, through spring to summer (Kruskal-Wallis $X^2$=13.01, d.f.=2, P=0.001). Median within-nest relatedness decreased non-significantly from winter to BPP summer (2-way ordered ANOVA by season and year: $F_{(3,123)}$=1.70, P=0.17). NPP = Nestmate provisioning phase, BPP = Brood provisioning phase.

| **Colony phase** | **Proportion of nests that contain full sisters** | | | **Within nest relatedness** | | |
|---|---|---|---|---|---|---|
| | 2012 | 2013 | **Mean** | 2012 | 2013 | **Mean** |
| Winter | 0.41±0.29 (14) | NA | **0.41±0.29** | 0.35±0.21 (14) | NA | **0.35±0.21** |
| NPP | 0.20±0.33 (35) | 0.30±0.43 (24) | **0.24±0.38** | 0.24±0.29 (35) | 0.18±0.40 (24) | **0.21±0.34** |
| BPP | 0.20±0.38 (41) | 0.17±0.37 (20) | **0.19±0.37** | 0.19±0.33 (41) | 0.09±0.42 (20) | **0.16±0.36** |
| 2012 vs. 2013 | Kruskal-Wallis $X^2$=13.01, d.f.=2, P=0.001 | | | 2-way ordered ANOVA by season and year: $F_{(3,123)}$=1.70, P=0.17 | | |

**Supplementary Table 3.** Microsatellite genotypes for all Xylocopa virginica females captured in 2012 and 2013 that were used for inferring sibling relationships and relatedness.

| Year | Individual ID | Nest Foraged In | XV23 | XV39 | XV42 | XV43 | XV3 | XV27 | XV7 | XV24 | XV30 |
|---|---|---|---|---|---|---|---|---|---|---|---|
| 2012 | A-10-3 | A-10 | 383/392 | 237/237 | 459/459 | 210/213 | 219/219 | 218/230 | 316/316 | 198/198 | 288/298 |
| 2012 | A-U-8 | A-10 | 383/383 | 237/240 | 453/459 | 204/210 | 223/223 | 221/221 | 316/316 | 189/204 | 298/298 |
| 2012 | B-U-9 | A-10 | 392/392 | 240/240 | 453/453 | / | 219/235 | 218/221 | 316/316 | 198/198 | 288/295 |
| 2012 | A-11-2 | A-11 | 392/392 | 240/240 | 459/462 | 210/210 | 219/219 | 218/221 | 316/316 | 189/198 | 288/298 |
| 2012 | A-11-3 | A-11 | 380/412 | 240/240 | 453/459 | 207/207 | 219/219 | 218/230 | 316/316 | 189/198 | 298/301 |
| 2012 | A-11-6 | A-11 | 392/392 | 240/240 | 453/453 | 204/207 | 219/235 | 218/230 | 316/316 | 189/198 | 295/301 |
| 2012 | A-12-1 | A-12 | 383/392 | 240/240 | 459/459 | 210/210 | 219/235 | 218/218 | 316/316 | 198/198 | 288/295 |
| 2012 | A-12-2 | A-12 | 383/389 | 240/240 | 459/459 | 207/210 | 219/235 | 218/218 | 316/316 | 189/198 | 288/295 |
| 2012 | A-U-15 | A-13 | 383/383 | 237/240 | 453/453 | 204/210 | 219/219 | 218/221 | 316/316 | 198/201 | 288/298 |
| 2012 | C-2B-1 | A-14 | 380/380 | 240/240 | 453/459 | 210/219 | 219/223 | 230/239 | 316/320 | 204/204 | 298/298 |
| 2012 | B-U-4 | A-15 | 380/412 | 240/240 | 453/459 | 210/210 | 219/219 | 218/230 | 316/316 | 198/198 | 298/298 |
| 2012 | A-U-9 | A-16 | 383/383 | 237/240 | 453/459 | 204/210 | 223/223 | 221/221 | 316/316 | 189/204 | 298/298 |
| 2012 | A-U-17/A-16 | A-16 | 383/383 | 237/237 | 453/459 | 210/213 | 219/219 | 221/221 | 316/316 | 189/198 | 298/310 |
| 2012 | A-2-1 | A-2 | 383/383 | 237/240 | 453/453 | 207/210 | 223/223 | 218/230 | 316/316 | 201/204 | 298/298 |
| 2012 | A-2-5 | A-2 | 383/383 | 240/240 | 453/453 | 207/210 | 219/223 | 218/230 | 316/316 | 201/204 | 288/298 |
| 2012 | A-3-3 | A-3 | 392/412 | 240/240 | 459/459 | 207/210 | 219/219 | 221/221 | 316/316 | 198/198 | 298/298 |
| 2012 | A-U-11 | A-3 | 389/412 | 240/240 | 453/459 | 210/210 | 219/219 | 221/221 | 316/316 | 198/198 | 298/298 |
| 2012 | A-4-2 | A-4 | 380/383 | 240/240 | 453/453 | 207/210 | 219/219 | 221/230 | 316/320 | 195/198 | 298/298 |
| 2012 | A-4-1 | A-4 | 380/392 | 240/240 | 453/453 | 207/207 | 219/219 | 218/230 | 316/320 | 195/198 | 298/298 |
| 2012 | A-6-1 | A-6 | 392/398 | 237/240 | 462/464 | 210/210 | 223/223 | 227/239 | 316/320 | 195/198 | 298/298 |
| 2012 | A-6-2 | A-6 | 389/412 | 237/240 | 462/464 | 210/213 | 223/223 | 230/239 | 316/320 | 198/198 | 298/298 |
| 2012 | A-2-6 | A-7 | 380/383 | 240/240 | 453/459 | 207/213 | 223/223 | 218/233 | 316/316 | 198/204 | 298/298 |

| | | | | | | | | | | | |
|---|---|---|---|---|---|---|---|---|---|---|---|
| 2012 | A-9-3 | A-7 | 380/392 | 240/240 | 453/459 | 207/210 | 219/219 | 218/224 | 316/316 | 198/204 | 298/310 |
| 2012 | A-8-15 | A-8 | 380/383 | 237/237 | 459/462 | 210/210 | 219/223 | 218/230 | 316/316 | 198/204 | 288/298 |
| 2012 | A-U-16 | A-8 | 380/380 | 237/240 | 453/462 | 204/210 | 219/223 | 218/239 | 316/316 | 192/204 | 288/298 |
| 2012 | A-8-9 | A-9 | 380/383 | 237/240 | 453/462 | 210/210 | 219/223 | 218/230 | 316/316 | 198/204 | 298/298 |
| 2012 | A-9-1 | A-9 | 392/392 | 237/240 | 453/456 | 210/210 | 219/247 | 221/233 | 316/316 | 198/198 | 298/301 |
| 2012 | A-9-6 | A-9 | 380/383 | 237/240 | 453/453 | 207/210 | 223/223 | 230/236 | 316/316 | 195/198 | 298/298 |
| 2012 | B-U-7 | B-1 | 383/389 | 237/240 | 453/453 | 210/210 | 219/219 | / | 316/316 | 198/198 | 298/298 |
| 2012 | C-3-1 | B-1 | 380/392 | 240/240 | / | 207/213 | 219/223 | 221/230 | 316/320 | 198/201 | 298/298 |
| 2012 | A-8-13 | B-10 | 380/383 | 237/237 | 453/462 | 204/210 | 219/223 | 230/239 | 316/316 | 192/198 | 288/298 |
| 2012 | B-2-2 | B-2 | 380/412 | 240/240 | 459/459 | / | 219/219 | 221/221 | 316/316 | 198/204 | / |
| 2012 | B-2-3 | B-2 | 383/392 | 237/240 | 453/459 | 204/210 | 223/223 | 221/233 | 316/316 | 198/198 | 298/298 |
| 2012 | B-3-4 | B-3 | 383/389 | 240/240 | 453/453 | 204/207 | 219/232 | 227/230 | 320/320 | 189/198 | 288/310 |
| 2012 | B-4-1 | B-4 | 392/412 | / | / | 210/210 | 219/219 | 221/221 | 316/316 | 198/198 | 298/298 |
| 2012 | B-4-3 | B-4 | 383/389 | 240/240 | 453/453 | / | 219/219 | 227/233 | 316/320 | 198/198 | 288/298 |
| 2012 | B-5-2 | B-5 | 383/386 | 237/240 | 453/459 | 207/210 | 219/223 | 221/221 | 316/316 | 189/204 | 298/298 |
| 2012 | B-6-2 | B-6 | 380/392 | 240/240 | 453/453 | 210/210 | 219/227 | 221/221 | 316/316 | 189/198 | 298/298 |
| 2012 | B-6-3 | B-6 | 380/392 | 240/240 | 453/459 | 210/210 | 219/227 | 218/227 | 316/316 | 198/198 | 298/298 |
| 2012 | B-7-1 | B-7 | 380/392 | 237/240 | 453/453 | 210/210 | 219/223 | 221/221 | 316/316 | 198/198 | 298/298 |
| 2012 | A-11-5 | B-8 | 392/392 | 237/240 | 453/453 | 204/210 | 235/235 | 218/221 | 316/316 | 198/204 | 295/298 |
| 2012 | D-U-23 | B-8 | 383/389 | 240/240 | 462/464 | 210/213 | 219/219 | 218/221 | 316/316 | 198/204 | 298/298 |
| 2012 | A-9-5 | B-9 | 380/383 | 237/240 | 453/453 | 207/210 | 223/223 | 218/230 | 316/320 | 195/198 | 298/298 |
| 2012 | B-10-1 | B-9 | 380/412 | 240/240 | / | 210/210 | 219/219 | / | 316/316 | 201/204 | 298/298 |
| 2012 | A-U-7 | C-1 | 392/412 | 237/240 | 453/459 | / | 219/219 | 218/218 | 316/316 | 198/198 | 298/298 |
| 2012 | C-1-1 | C-1 | 383/383 | 240/243 | / | 207/210 | 223/223 | 221/230 | 316/316 | 198/198 | 298/298 |
| 2012 | C-1-2 | C-1 | 380/383 | 240/243 | 453/453 | 210/210 | 219/219 | 218/221 | 316/320 | 198/198 | 298/298 |
| 2012 | C-2-2 | C-2 | 383/392 | 240/240 | 453/453 | 204/213 | 219/219 | 218/221 | 316/316 | 198/204 | 298/298 |
| 2012 | C-3-2 | C-3 | 383/395 | 240/240 | 453/459 | 210/210 | 219/219 | 221/230 | 316/316 | 198/204 | 298/298 |
| 2012 | C-4-1 | C-4 | 380/383 | 237/240 | 453/453 | 207/207 | 219/223 | 218/230 | 316/320 | 198/204 | 298/298 |

| | | | | | | | | | | | |
|---|---|---|---|---|---|---|---|---|---|---|---|
| 2012 | C-4-2 | C-4 | 380/398 | 237/240 | 453/464 | 207/210 | 223/223 | 224/230 | 316/316 | 189/198 | 298/298 |
| 2012 | B-8-1 | C-4 | 380/389 | 240/240 | 453/464 | 213/213 | 219/223 | 221/221 | 316/316 | 198/201 | 298/298 |
| 2012 | C-5-1 | C-5 | 380/383 | 237/240 | 453/456 | 207/210 | 219/240 | 230/233 | 316/316 | 198/198 | 298/310 |
| 2012 | C-5-2 | C-5 | 383/412 | 237/237 | 450/450 | 210/210 | 219/223 | 230/233 | 316/316 | 189/198 | 298/310 |
| 2012 | A-11-4 | C-6 | 380/386 | 240/240 | 453/462 | 207/207 | 219/219 | 218/221 | 320/320 | 198/198 | 301/310 |
| 2012 | A-9-2 | C-6 | 392/392 | 237/240 | 456/459 | 210/210 | 219/247 | 221/233 | 316/316 | 198/198 | 298/301 |
| 2012 | C-7-1 | C-6 | / | 237/240 | 456/456 | 207/213 | 219/219 | 218/230 | 316/320 | / | 298/298 |
| 2012 | C-8-1 | C-8 | 380/392 | 240/240 | 453/453 | 204/207 | 223/223 | 218/230 | 316/316 | 198/204 | 295/298 |
| 2012 | C-8-4 | C-8 | 383/412 | 237/240 | 453/453 | 207/210 | 223/223 | 230/230 | 316/320 | 198/204 | 298/298 |
| 2012 | D-1-1 | D-1 | 380/383 | 240/240 | 453/453 | 210/210 | 219/219 | 230/230 | 316/320 | 198/204 | 288/288 |
| 2012 | D-1-4 | D-1 | 383/383 | 240/240 | 462/462 | 219/219 | 240/240 | 218/221 | 316/316 | 198/198 | 298/298 |
| 2012 | D-1-6 | D-1 | 383/383 | 237/240 | 453/453 | 210/219 | 219/232 | 218/230 | 316/316 | 204/204 | 298/298 |
| 2012 | D-U-16 | D-11 | 383/389 | 237/240 | 453/462 | 207/210 | 219/232 | 218/230 | 316/320 | 198/201 | 298/298 |
| 2012 | D-U-20 | D-11 | 383/383 | 237/237 | 453/462 | 207/207 | 232/240 | 218/230 | 316/320 | 198/201 | 298/298 |
| 2012 | D-12-1 | D-12 | 380/392 | 237/240 | 453/459 | 207/207 | 219/219 | 221/221 | 316/320 | 198/198 | 298/298 |
| 2012 | D-12-2 | D-12 | 380/383 | 240/240 | 459/462 | / | 219/219 | 218/230 | 316/316 | 198/204 | 298/298 |
| 2012 | D-12-3 | D-12 | 383/392 | 240/240 | 453/453 | 207/210 | 219/232 | 221/230 | 316/316 | 198/204 | 298/298 |
| 2012 | D-U-1 | D-14 | 395/395 | 240/240 | 462/464 | / | 232/232 | 218/221 | 316/316 | 189/189 | 298/298 |
| 2012 | D-U-12 | D-14 | 383/392 | 240/240 | 453/464 | 207/207 | 232/232 | 221/230 | 316/316 | 198/204 | 298/298 |
| 2012 | D-15-1 | D-15 | 383/392 | 240/240 | 453/462 | 210/210 | 219/219 | 230/230 | 316/316 | 198/198 | 288/298 |
| 2012 | D-U-11 | D-15 | 380/392 | 237/237 | 459/464 | 207/207 | 223/232 | 218/227 | 316/316 | 189/204 | 298/298 |
| 2012 | D-U-6 | D-15 | 392/398 | / | 459/462 | 204/210 | 219/240 | 218/218 | 316/316 | 198/198 | 298/298 |
| 2012 | D-U-5 | D-16 | 380/392 | 237/237 | 453/453 | 207/207 | 219/219 | 221/221 | 316/320 | 198/198 | 301/301 |
| 2012 | D-U-21 | D-17 | 380/392 | 237/240 | 453/453 | 204/207 | 219/219 | 221/230 | 316/320 | 198/198 | 298/298 |
| 2012 | D-U-8 | D-17 | 380/392 | 237/240 | 453/453 | 204/207 | 219/219 | 230/230 | 316/320 | 198/198 | 298/298 |
| 2012 | D-19-1 | D-19 | 380/383 | 237/240 | 453/459 | 204/210 | 223/223 | 218/230 | 316/320 | 198/198 | 298/298 |
| 2012 | D-2-4 | D-2 | 383/395 | 240/240 | / | 204/210 | 219/219 | 218/230 | 316/320 | 198/198 | 288/288 |
| 2012 | D-U-3 | D-2 | 380/392 | / | 456/456 | 204/210 | 219/219 | / | 316/316 | / | 298/298 |

| | | | | | | | | | | | |
|---|---|---|---|---|---|---|---|---|---|---|---|
| 2012 | D-8-5 | D-2 | 380/383 | 237/240 | 453/453 | 204/210 | 219/232 | 230/230 | 316/316 | 198/198 | 288/298 |
| 2012 | D-5-2 | D-20 | 383/412 | 237/240 | 453/464 | 204/207 | 223/223 | 218/221 | 316/316 | 198/201 | 298/298 |
| 2012 | D-3-3 | D-3 | 380/392 | 237/240 | 459/464 | 210/210 | 219/240 | 218/218 | 316/324 | 198/198 | 298/298 |
| 2012 | D-3-5 | D-3 | 383/398 | 240/240 | 453/464 | 210/210 | 219/219 | 218/230 | 316/324 | 198/198 | 298/298 |
| 2012 | D-3-6 | D-3 | 380/383 | 237/240 | 453/453 | 204/210 | 232/240 | 218/230 | 316/324 | 198/198 | 298/298 |
| 2012 | D-4-6 | D-4 | 383/392 | 240/240 | 453/459 | 210/219 | 219/219 | 218/230 | 316/316 | 189/198 | 298/298 |
| 2012 | D-U-29 | D-4 | 383/383 | 240/240 | 453/453 | 204/210 | 219/219 | 221/230 | 316/316 | 198/204 | 288/298 |
| 2012 | D-U-22 | D-4 | 383/412 | 237/240 | 453/453 | 204/207 | 223/232 | 230/230 | 316/316 | 198/198 | 298/298 |
| 2012 | D-5-1 | D-5 | 383/383 | 237/240 | 453/462 | 210/210 | 232/240 | 218/218 | 316/316 | 198/198 | 298/298 |
| 2012 | D-U-7 | D-5 | 380/380 | 240/240 | 453/453 | 210/210 | 219/223 | 221/221 | 316/316 | 198/204 | 288/298 |
| 2012 | D-U-27 | D-5 | 383/412 | 240/240 | 453/464 | 204/204 | 223/223 | 218/221 | 316/320 | 198/201 | 298/298 |
| 2012 | D-U-18 | D-5 | 383/412 | 240/240 | 453/464 | 204/204 | 219/223 | 218/221 | 316/320 | 198/204 | 288/298 |
| 2012 | D-U-14 | D-6 | 383/389 | 237/237 | 453/462 | 207/207 | 240/240 | 218/230 | 316/320 | 198/201 | 298/298 |
| 2012 | D-8-3 | D-8 | 380/383 | 237/240 | 450/450 | 204/207 | 219/219 | 230/230 | 316/316 | 198/198 | 288/298 |
| 2012 | D-8-4 | D-8 | 380/383 | 237/240 | 453/453 | 204/207 | 219/219 | 230/230 | 316/316 | 198/204 | 288/298 |
| 2012 | D-8-6 | D-8 | 380/383 | 237/240 | 453/453 | 204/210 | 232/232 | 230/230 | 316/320 | 195/198 | 298/298 |
| 2012 | D-9-10B | D-9 | 383/383 | 237/240 | 453/459 | 210/210 | 219/219 | 227/227 | 316/316 | 189/198 | 288/298 |
| 2012 | F-1-1 | F-1 | 380/380 | 237/240 | 453/459 | 207/207 | 219/223 | 218/221 | 316/320 | 204/204 | 298/301 |
| 2012 | F-1-3 | F-1 | 383/392 | 240/240 | 464/464 | 204/210 | 219/219 | 218/233 | 316/320 | 195/195 | 298/298 |
| 2012 | F-U-15 | F-1 | 383/392 | 240/240 | 453/464 | 204/207 | 223/232 | 218/233 | 316/316 | 195/198 | 298/298 |
| 2012 | B-3B-1 | F-10 | 380/380 | / | 453/453 | 207/210 | 219/219 | 230/230 | 316/316 | 204/204 | 298/310 |
| 2012 | F-U-17 | F-11 | 380/380 | 240/240 | 453/462 | 204/210 | 232/240 | 218/224 | 308/316 | 189/198 | 298/298 |
| 2012 | C-2B-5 | F-11 | 383/383 | 237/237 | 453/462 | 207/210 | 219/223 | 224/239 | 316/316 | 198/198 | 298/298 |
| 2012 | F-16-2 | F-12 | 383/389 | 237/237 | 453/453 | 204/207 | 232/232 | 224/233 | 316/316 | 189/198 | 298/298 |
| 2012 | F-13-1 | F-14 | 380/383 | 237/240 | 453/453 | 207/210 | 232/232 | 218/224 | 308/316 | 198/198 | 298/310 |
| 2012 | F-14-1 | F-14 | 383/389 | 240/240 | 453/462 | 210/213 | 223/223 | 218/221 | 316/320 | 198/204 | 298/298 |
| 2012 | F-15-1 | F-15 | 380/383 | 237/240 | / | 210/210 | 219/227 | 218/230 | 316/320 | 198/198 | 288/310 |
| 2012 | F-15-2 | F-15 | 380/383 | / | / | 210/210 | 232/232 | 218/227 | 316/316 | 198/198 | 298/298 |

| | | | | | | | | | | | |
|---|---|---|---|---|---|---|---|---|---|---|---|
| 2012 | F-U-13 | F-15 | 380/383 | 237/240 | 453/462 | 210/210 | 219/232 | 218/227 | 316/320 | 198/198 | 288/288 |
| 2012 | F-16-1 | F-16 | 383/392 | 240/240 | 453/459 | 204/210 | 232/232 | 227/233 | 316/316 | 198/204 | 298/298 |
| 2012 | F-U-11 | F-16 | 380/412 | 240/243 | 453/459 | 210/210 | 223/223 | 221/230 | 316/316 | 189/189 | 298/301 |
| 2012 | F-17-1 | F-17 | 383/389 | 237/240 | 453/453 | 207/210 | 219/219 | 218/233 | 316/316 | 189/198 | 298/301 |
| 2012 | D-6A-2 | F-18 | 380/383 | 240/240 | 450/462 | 210/210 | 219/219 | 230/233 | 316/316 | 189/204 | 298/298 |
| 2012 | F-2-1 | F-2 | 380/380 | / | / | 210/210 | 223/240 | 218/230 | 316/320 | 189/198 | 298/301 |
| 2012 | F-20-1 | F-20 | 383/392 | 240/243 | 453/464 | 210/210 | 219/219 | 224/230 | 316/316 | 195/198 | 288/298 |
| 2012 | F-7-3 | F-22 | 392/412 | / | 462/462 | 207/210 | 219/232 | 218/227 | 316/316 | 195/198 | 301/310 |
| 2012 | F-3-1 | F-3 | 380/412 | 237/240 | / | 204/210 | 232/232 | 230/230 | 316/316 | 195/201 | 298/298 |
| 2012 | F-3-3 | F-3 | 383/383 | 240/243 | / | 213/213 | 219/219 | 221/236 | 316/316 | 198/201 | 298/301 |
| 2012 | F-4-1 | F-4 | 380/383 | 240/240 | 453/453 | 207/210 | 219/232 | 218/218 | 316/316 | 198/204 | 310/310 |
| 2012 | F-7-2 | F-4 | 380/380 | 240/240 | 453/459 | 204/213 | 219/223 | 218/221 | 316/316 | 198/204 | 298/301 |
| 2012 | F-6-2 | F-6 | 380/383 | 237/240 | 453/464 | 204/210 | 232/232 | 218/230 | 316/320 | 189/198 | 298/298 |
| 2012 | F-6-3 | F-6 | 380/383 | 237/240 | 453/459 | 210/213 | 219/219 | 218/230 | 316/320 | 189/198 | 298/298 |
| 2012 | F-U-10 | F-7 | 392/412 | 240/240 | 462/462 | 207/210 | 219/232 | 218/227 | 316/316 | / | 301/310 |
| 2013 | A-10 | A-10-2 | 383/392 | 237/237 | 453/459 | 210/213 | 219/219 | 218/230 | 316/316 | 198/198 | 288/288 |
| 2013 | A-10 | A-10-1 | 380/383 | 237/237 | 453/459 | 210/213 | 219/219 | 218/230 | 316/316 | 198/198 | 288/288 |
| 2013 | A-11 | A-11-3 | 389/392 | / | / | / | 236/236 | 218/221 | 316/316 | 198/198 | 298/301 |
| 2013 | A-11 | A-11-2 | 383/392 | 240/240 | 459/459 | 207/210 | 219/219 | 218/230 | 316/316 | 189/198 | 295/301 |
| 2013 | A-12 | A-9-1 | 380/383 | 237/240 | 453/453 | 207/210 | 223/223 | 230/236 | 316/316 | 195/198 | 298/298 |
| 2013 | A-13 | A-13-2 | 383/383 | 237/237 | 453/459 | 204/210 | 223/223 | 221/230 | 316/316 | 198/204 | / |
| 2013 | A-14 | A-14-4 | 380/380 | 237/240 | 453/459 | 204/213 | 219/219 | 218/233 | 316/320 | 198/204 | 298/310 |
| 2013 | A-14 | A-14-2 | 380/383 | 237/240 | 453/453 | / | 219/232 | 230/230 | 316/320 | 198/198 | 298/298 |
| 2013 | A-15 | A-15-1 | 380/383 | 237/237 | 459/459 | 210/213 | 219/219 | 221/230 | 316/316 | 198/204 | 288/298 |
| 2013 | A-16 | A-16-1 | 383/383 | 237/240 | 453/462 | 210/210 | 219/219 | 221/230 | 316/316 | 198/198 | 298/310 |
| 2013 | A-16 | A-7-2 | 380/392 | / | 453/459 | 207/210 | 219/223 | 230/233 | 320/324 | / | / |
| 2013 | A-17 | A-3-2 | 389/412 | 240/240 | 459/459 | 210/210 | 219/219 | 221/221 | 316/316 | 198/198 | 298/298 |
| 2013 | A-18 | A-4-2 | 380/383 | 237/240 | 459/462 | 210/210 | 219/223 | 218/230 | 316/316 | 198/204 | 288/298 |

| | | | | | | | | | | | |
|---|---|---|---|---|---|---|---|---|---|---|---|
| 2013 | A-2 | A-2-1 | 380/383 | 240/240 | 453/459 | 207/213 | 223/223 | 230/233 | 316/316 | 198/204 | 298/298 |
| 2013 | A-2 | A-2-2 | 383/398 | 237/237 | 453/453 | 204/207 | 223/223 | 230/230 | 316/316 | 198/201 | 298/298 |
| 2013 | A-3 | D-5A-1 | 383/383 | 237/240 | 453/462 | 210/210 | 219/224 | 230/233 | 316/316 | 198/198 | 298/298 |
| 2013 | A-4 | A-4-1 | 383/383 | 237/237 | 450/453 | 210/210 | 223/223 | 221/227 | 316/316 | 198/198 | 288/298 |
| 2013 | A-6 | A-6-1 | 380/392 | 237/237 | 453/459 | / | 223/223 | 218/239 | 316/320 | 198/198 | 298/301 |
| 2013 | A-7 | A-7-1 | 380/392 | 240/240 | 453/459 | 207/210 | 219/223 | 230/233 | 316/320 | 198/198 | 298/298 |
| 2013 | A-8 | A-8-1 | 383/383 | 237/240 | 459/462 | 210/210 | 219/219 | 218/230 | 316/316 | 198/198 | 288/298 |
| 2013 | A-8 | A-8-2 | 380/383 | 237/240 | 453/453 | 210/213 | 219/219 | 218/221 | 316/316 | 198/198 | 298/310 |
| 2013 | A-9 | A-9-2 | 380/386 | 237/237 | 453/459 | 204/210 | 219/219 | 221/230 | 316/316 | 195/198 | 298/298 |
| 2013 | B-10 | B-10-1 | 380/380 | 237/237 | 462/462 | 210/210 | 219/223 | 224/239 | 316/316 | 192/195 | 288/298 |
| 2013 | B-10 | B-10-2 | 380/383 | 237/237 | 453/462 | 204/210 | 223/223 | 224/239 | 316/316 | 192/195 | 298/298 |
| 2013 | B-2 | B-2-1 | 380/392 | 237/240 | 453/462 | / | 223/223 | 218/239 | 316/316 | 198/198 | 298/298 |
| 2013 | B-2 | B-2-2 | 383/392 | 240/240 | 453/453 | 204/207 | 223/223 | 218/230 | 316/316 | 198/204 | 295/298 |
| 2013 | B-5 | B-5-2 | 380/383 | 237/240 | 453/453 | 210/210 | 223/223 | 218/230 | 316/320 | 195/198 | 298/298 |
| 2013 | B-5 | C-7-1 | 383/383 | 237/237 | 453/459 | 210/210 | 223/223 | 224/233 | 316/316 | 198/204 | 298/298 |
| 2013 | B-7 | B-7-3 | 383/392 | 237/240 | 453/459 | 204/210 | 219/232 | 221/233 | 316/320 | 198/198 | 298/310 |
| 2013 | B-7 | B-7-4 | 380/383 | 237/240 | 453/462 | 210/210 | 219/219 | 230/230 | 316/316 | 198/204 | 298/298 |
| 2013 | B-7 | B-7-2 | 383/392 | 237/240 | 453/459 | 204/213 | 223/223 | 221/233 | 316/320 | 198/198 | 298/298 |
| 2013 | B-8 | B-8-1 | 383/392 | 240/240 | 459/462 | 204/210 | 219/219 | 218/230 | 316/320 | 195/195 | 298/310 |
| 2013 | B-9 | B-9-1 | 380/383 | 237/240 | 453/456 | 210/210 | 219/223 | 230/230 | 316/316 | 189/198 | 298/298 |
| 2013 | C-2 | C-2-1 | 380/392 | 240/240 | 453/453 | 204/207 | 223/236 | 218/230 | 316/316 | 192/198 | 295/298 |
| 2013 | C-5 | C-5-1 | 383/383 | 237/243 | 453/453 | 210/213 | 223/223 | 230/230 | 316/320 | 189/198 | 298/298 |
| 2013 | C-5 | C-5-3 | 383/383 | 237/237 | 453/456 | 204/207 | 219/219 | 230/230 | 316/320 | 198/198 | 298/298 |
| 2013 | C-5 | C-5-2 | 383/383 | 240/240 | 453/459 | 210/210 | 240/240 | 218/224 | 316/320 | 198/204 | 298/298 |
| 2013 | D-1 | D-1-1 | 380/383 | 240/243 | 453/453 | 207/210 | 219/219 | 230/230 | 316/316 | 195/195 | 288/288 |
| 2013 | D-1 | D-1-3 | 380/395 | 237/240 | 453/462 | 204/210 | 232/232 | 221/221 | 316/320 | 198/198 | 298/298 |
| 2013 | D-11 | D-11-1 | 380/383 | 240/240 | 459/459 | 219/219 | 219/219 | 221/230 | 316/316 | 189/198 | 298/298 |
| 2013 | D-11 | D-11-4 | 380/383 | 237/240 | 459/462 | 207/210 | 232/240 | 218/221 | 316/316 | 198/198 | 298/298 |

| | | | | | | | | | | | |
|---|---|---|---|---|---|---|---|---|---|---|---|
| 2013 | D-12 | D-11-3 | 383/383 | 240/240 | 453/453 | 207/210 | 219/219 | 218/230 | 316/316 | 198/198 | 298/298 |
| 2013 | D-12 | D-12-2 | 383/389 | 237/240 | 453/462 | 210/213 | 219/219 | 218/230 | 316/316 | 195/198 | 298/301 |
| 2013 | D-13 | D-13-1 | 380/383 | 237/237 | 453/453 | 207/210 | 219/219 | / | 316/316 | 198/204 | 288/298 |
| 2013 | D-14 | D-14-1 | 380/383 | / | / | 207/207 | / | 221/230 | 316/320 | 198/201 | 298/298 |
| 2013 | D-15 | D-15-1 | 383/398 | 240/240 | 453/462 | 204/210 | 219/219 | 218/230 | 316/316 | 198/198 | 288/298 |
| 2013 | D-16 | D-16-1 | 392/392 | 240/243 | 453/464 | 207/207 | 219/223 | 218/224 | 316/316 | 198/198 | 298/301 |
| 2013 | D-19 | D-19-1 | 380/398 | 240/240 | 453/462 | 210/210 | 219/219 | 230/230 | 316/316 | 198/198 | 288/288 |
| 2013 | D-19 | D-8-2 | 380/383 | 237/240 | 453/453 | 204/204 | 219/223 | 230/230 | 316/316 | 198/198 | 288/298 |
| 2013 | D-2 | D-2-2 | 380/392 | 240/240 | 453/459 | 207/210 | 232/232 | / | 316/316 | 198/198 | 298/298 |
| 2013 | D-20 | D-20-1 | 383/383 | 237/240 | 453/453 | 210/219 | 232/232 | 218/230 | 316/316 | 204/204 | 298/301 |
| 2013 | D-3 | D-3-1 | 383/383 | 240/240 | 453/464 | 210/210 | 219/219 | 218/230 | 316/324 | 198/201 | 288/298 |
| 2013 | D-3 | D-3-3 | 380/383 | 240/240 | 453/464 | 210/210 | 219/219 | 218/221 | 316/316 | 198/201 | 288/298 |
| 2013 | D-4 | D-4-1 | 383/383 | 240/240 | 453/462 | 207/210 | 219/219 | 221/230 | 316/316 | 198/198 | 288/298 |
| 2013 | D-4 | D-4-2 | 380/383 | 237/237 | 453/453 | 207/210 | 223/223 | / | 316/316 | 198/198 | 298/298 |
| 2013 | D-5 | D-6-1 | 380/403 | 237/240 | 453/459 | 210/213 | 219/219 | 218/221 | 316/316 | 198/198 | 298/298 |
| 2013 | D-6 | D-19-2 | 380/383 | 240/240 | 453/459 | 207/207 | 240/240 | 218/221 | 316/320 | 198/198 | 298/298 |
| 2013 | D-8 | D-8-1 | 383/389 | 237/240 | 453/453 | 204/207 | 219/223 | 230/230 | 316/316 | 198/198 | 288/298 |
| 2013 | D-8 | D-8-3 | 383/383 | 240/243 | 453/462 | 207/213 | 232/232 | 221/230 | 316/316 | 198/204 | 298/298 |
| 2013 | D-9 | D-1-2 | 383/392 | 237/240 | 453/462 | 204/207 | 223/223 | 221/230 | 316/316 | 198/198 | 298/298 |
| 2013 | D-9 | D-9-1 | 383/383 | 237/240 | 453/453 | 207/210 | 219/223 | 230/230 | 320/324 | 189/204 | 298/298 |
| 2013 | D-9 | D-9-3 | 383/383 | 237/240 | 453/459 | 207/210 | 219/240 | 218/230 | 316/320 | 189/204 | 298/310 |
| 2013 | F-1 | F-1-1 | 383/395 | 240/240 | 453/462 | 204/210 | / | 218/233 | 316/316 | 195/198 | 298/298 |
| 2013 | F-1 | F-1-2 | 380/383 | 237/237 | 453/459 | 207/213 | 219/219 | 221/230 | 316/320 | 198/204 | 298/301 |
| 2013 | F-10 | F-10-1 | 380/380 | 240/240 | 453/453 | 207/210 | 232/232 | 218/224 | 316/316 | 198/198 | 298/298 |
| 2013 | F-12 | F-4-2 | 380/380 | 240/240 | 453/462 | 207/210 | 219/219 | 218/230 | 316/316 | 198/204 | / |
| 2013 | F-15 | F-15-1 | 380/383 | 237/240 | 459/462 | 210/210 | 219/219 | 218/230 | 316/316 | 198/198 | 288/288 |
| 2013 | F-2 | F-2-1 | 380/383 | 240/243 | 459/464 | 207/210 | 219/223 | 221/230 | 316/316 | 189/198 | 298/298 |
| 2013 | F-20 | F-7-1 | 380/412 | 240/240 | 462/464 | 207/210 | 219/219 | 227/230 | 316/316 | 198/204 | 298/301 |

| | | | | | | | | | | | |
|---|---|---|---|---|---|---|---|---|---|---|---|
| 2013 | F-22 | D-5A-4 | 383/412 | 240/240 | 448/448 | / | 224/224 | 230/230 | 316/320 | 204/204 | 298/298 |
| 2013 | F-3 | F-3-1 | 383/383 | 237/240 | 453/462 | 210/210 | 219/219 | 221/230 | 316/316 | 204/204 | 298/298 |
| 2013 | F-3 | F-3-3 | 380/383 | 240/240 | 453/459 | 204/210 | 219/219 | 218/218 | 316/320 | 198/198 | / |
| 2013 | F-4 | F-4-1 | 380/383 | 240/240 | 453/462 | 204/210 | 219/219 | 218/230 | 316/316 | 198/204 | 298/298 |
| 2013 | F-6 | F-3-2 | 398/412 | 237/240 | 453/453 | 204/210 | 223/232 | 218/230 | 316/316 | 195/198 | 298/298 |
| 2013 | F-7 | F-7-2 | 380/383 | / | 453/464 | 210/213 | 219/219 | 218/230 | 316/320 | 189/198 | 298/298 |

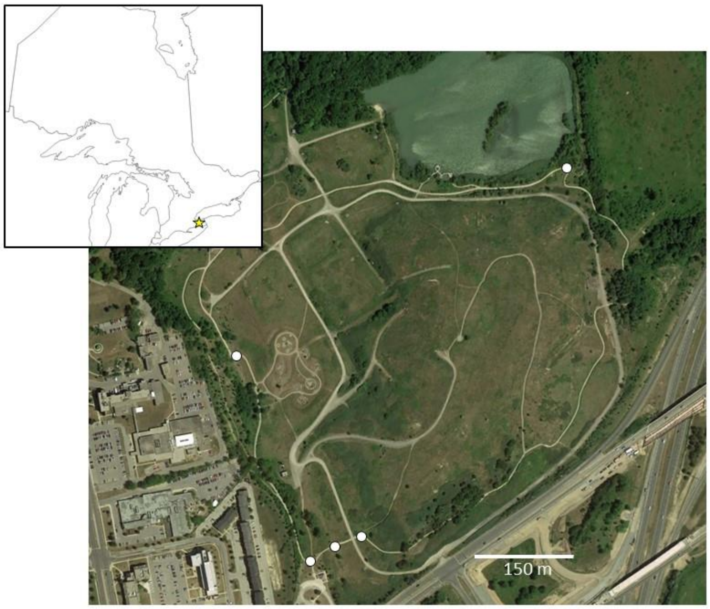


**Supplementary Figure 1**. Map of sample sites. Inset map indicates general location of field sites in southern Ontario, Canada. Colour map is an aerial view of the Glenridge Quarry Naturalization site in which all of our observations were made. The five white circles indicate the location of the 5 wooden bridges which were used by eastern carpenter bee females as nesting sites in 2012 and 2013.

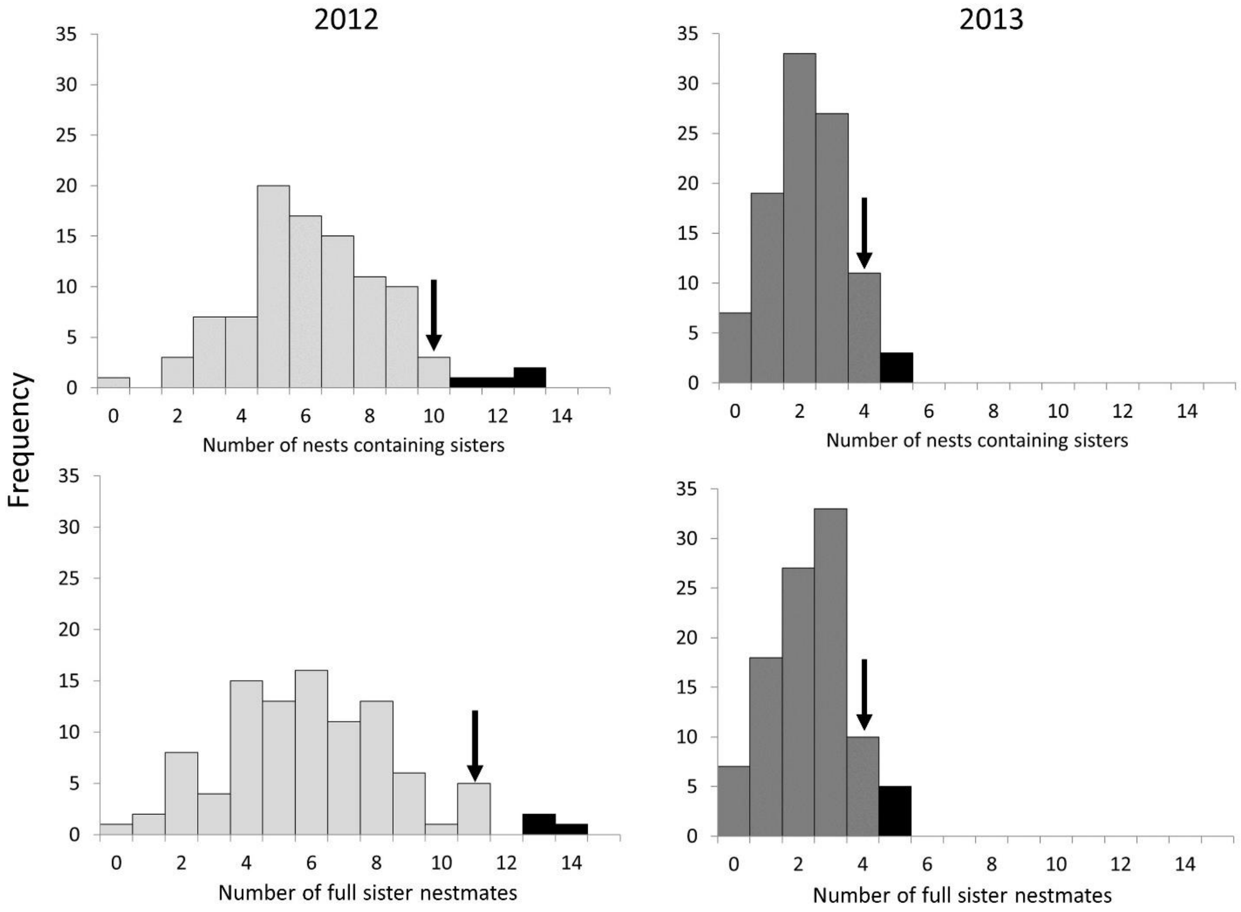


**Supplementary Figure 2**. Simulations to assess the number of sibships that would be observed if genotyped females were randomly distributed among available social colonies in 2012 and 2013. Observed numbers of full sisters that were nestmates during the brood provisioning period (black arrow) were compared to the numbers expected if adult females were randomly distributed among nests. Histograms represent distributions generated by 100 simulations, in which genotyped adult females were randomly assigned to colonies, constrained by the observed colony size distribution for each year. Black bars include values within the upper 5% of each distribution. To generate the 2012 distributions, 189 adult females were randomized among 71 nests, and for 2013, 101 females were randomized among 64 nests. The distributions are wider for 2012, the high-density year, because there were more females in more nests that year, than in the 2013, when density was significantly lower.
